# RstA is a Major Regulator of *Clostridioides difficile* Toxin Production and Motility

**DOI:** 10.1101/416115

**Authors:** Adrianne N. Edwards, Brandon R. Anjuwon-Foster, Shonna M. McBride

**Affiliations:** Department of Microbiology and Immunology, Emory University School of Medicine, Emory Antibiotic Resistance Center, Atlanta, GA, USA; Department of Microbiology and Immunology, University of North Carolina at Chapel Hill School of Medicine, Chapel Hill, NC, USA; Laboratory of Molecular Biology - Cell Biology Section, National Cancer Institute of the National Institutes of Health, Bethesda, MD, USA

**Keywords:** *Clostridium difficile*, anaerobe, RNPP, RRNPP, toxin, TcdA, TcdB, motility, transcriptional regulator, helix-turn-helix (HTH) motif, sporulation, spore

## Abstract

*Clostridioides difficile* infection (CDI) is a toxin-mediated disease. Several factors have been identified that influence the production of the two major *C. difficile* toxins, TcdA and TcdB, but prior published evidence suggested that additional unknown factors were involved in toxin regulation. Previously, we identified a *C. difficile* regulator, RstA, that promotes sporulation and represses motility and toxin production. We observed that the predicted DNA-binding domain of RstA was required for RstA-dependent repression of toxin genes, motility genes and *rstA* transcription. In this study, we further investigated the regulation of toxin and motility gene expression by RstA. DNA pulldown assays confirmed that RstA directly binds the *rstA* promoter via the predicted DNA-binding domain. Through mutational analysis of the *rstA* promoter, we identified several nucleotides that are important for RstA-dependent transcriptional regulation. Further, we observed that RstA directly binds and regulates the promoters of the toxin genes, *tcdA* and *tcdB*, as well as the promoters for the *sigD* and *tcdR* genes, which encode regulators of toxin gene expression. Complementation analyses with the *Clostridium perfringens* RstA ortholog and a multi-species chimeric RstA protein revealed that the *C. difficile* C-terminal domain is required for RstA DNA-binding activity, suggesting that species-specific signaling controls RstA function. Our data demonstrate that RstA is a transcriptional repressor that autoregulates its own expression and directly inhibits transcription of the two toxin genes and two positive toxin regulators, thereby acting at multiple regulatory points to control toxin production.

**IMPORTANCE:** *Clostridioides difficile* is an anaerobic, gastrointestinal pathogen of humans and other mammals. *C. difficile* produces two major toxins, TcdA and TcdB, which cause the symptoms of the disease, and forms dormant endospores to survive the aerobic environment outside of the host. A recently discovered regulatory factor, RstA, inhibits toxin production and positively influences spore formation. Herein, we determine that RstA directly represses toxin gene expression and gene expression of two toxin gene activators, TcdR and SigD, creating a complex regulatory network to tightly control toxin production. In addition, the ability for RstA to bind DNA and repress toxin production requires the species-specific domain predicted to respond to small quorum-sensing peptides. This study provides a novel regulatory link between *C. difficile* sporulation and toxin production. Further, our data suggest that *C. difficile* toxin production is regulated through a direct sensing mechanism.

## INTRODUCTION

*Clostridioides difficile* infection (CDI) is a nosocomial and community-acquired gastrointestinal disease that affects individuals with dysbiotic gut microbiota, which commonly occurs after antibiotic treatment (1, 2). Clinical outcomes range from mild diarrhea to severe disease symptoms, including sepsis and death (1). The two glycosylating exotoxins, TcdA and TcdB, elicit CDI symptoms and are indispensable for *C. difficile* virulence (3). Environmental and intracellular signals, including nutrient availability and metabolic cues, strongly influence toxin production (4-7). There are numerous identified *C. difficile* factors that control toxin gene expression in response to these signals (8-12); however, the regulatory pathways and molecular mechanisms that directly control toxin gene expression are not fully understood (13).

Our previous work identified a novel regulator, RstA, which depresses *C. difficile* toxin production and motility (14). RstA inhibits transcription of the toxin genes, *tcdA* and *tcdB*, the toxin-specific sigma factor, *tcdR*, and the flagellar-specific sigma factor, *sigD*, which is essential for motility and directs *tcdR* expression (11, 12, 14-16). In addition to repressing motility and toxin production, RstA positively influences *C. difficile* spore formation, which is critical for the survival of the bacterium outside of the host and for transmission from host-to-host, indicating that RstA regulates diverse phenotypes important for *C. difficile* pathogenesis.

The predicted secondary structure of RstA reveals three apparent domains: an N-terminal conserved helix-turn-helix DNA-binding domain, followed by a series of multiple tetratricopeptide repeat (TPR) domains comprising a putative Spo0F-like protein-binding domain, and a C-terminal putative quorum-sensing-like domain (14). These characteristic features place RstA in the RRNPP (Rap/Rgg/NprR/PlcR/PrgX; formerly, RNPP) family of proteins. RRNPP proteins are prevalent in Gram-positive organisms and regulate competence, sporulation, toxin production and other important survival and virulence phenotypes (17-19). The DNA-binding or protein-binding activity of RRNPP proteins are controlled by the direct binding of small, quorum-sensing peptides (19). The precursor proteins encoding the quorum-sensing peptides are often adjacent to the regulatory RRNPP protein, and are translated, exported, processed and reinternalized at high cell densities (20-25). In addition, RRNPP proteins often autoregulate their own expression, as is observed for RstA (14). The presence of these conserved domains within RstA provides insight into how RstA may regulate *C. difficile* toxin production, motility and sporulation.

To better understand the regulatory impact RstA exerts on *C. difficile* toxin production and sporulation, we examined the function of the conserved DNA-binding domain. Our previous study had shown that the DNA-binding domain is required for RstA-dependent regulation of *rstA* expression and toxin gene expression, but is expendable for sporulation regulation. Here, we demonstrate that RstA directly binds to its promoter via an imperfect inverted repeat, and directly binds the *sigD* and the toxin genes promoters. Further, our data demonstrate that RstA and SigD independently control toxin expression, creating a multi-tiered regulatory pathway by which RstA represses toxin production. Finally, we show that the *C. perfringens rstA* ortholog does not complement toxin production or sporulation in a *C. difficile rstA* mutant. However, a chimeric RstA protein containing the *C. perfringens* DNA-binding domain and the *C. difficile* Spo0F-binding and quorum-sensing-binding domains restores both sporulation and toxin production, providing evidence that the ability to respond to species-specific signaling is necessary for RstA DNA-binding activity.

## MATERIALS AND METHODS

### Bacterial strains and growth conditions

The bacterial strains and plasmids used in the study are listed in **Table S1**. *C. difficile* strains were routinely cultured in BHIS or TY medium (pH 7.4) supplemented with 2-5 μg/ml thiamphenicol and/or 1 μg/ml nisin as needed (26). Overnight cultures of *C. difficile* were supplemented with 0.1% taurocholate and 0.2% fructose to promote spore germination and prevent sporulation, respectively, as indicated (26, 27). *C. difficile* strains were cultured in a 37°C anaerobic chamber with an atmosphere of 10% H_2_, 5% CO_2_ and 85% N_2_, as previously described (28). *Escherichia coli* strains were grown at 37°C in LB (29) with 100 μg/ml ampicillin and/or 20 μg/ml chloramphenicol as needed. Kanamycin (50 μg/ml) was used for counterselection against *E. coli* after conjugation with *C. difficile*, as previously described (30).

### Strain and plasmid construction and accession numbers

Oligonucleotides used in this study are listed in **Table S2**. Details of vector construction are described in the Supplemental Material (**Fig. S1)**. *C. difficile* strains 630 (Genbank no. NC_009089.1) and R20291 (Genbank no. FN545816.1), *Clostridium acetobutylicum* ATCC 824 (Genbank no. NC_003030.1), *Clostridium sordellii* ATCC 9714 (Genbank no. APWR00000000) and *Clostridium perfringens* S13 (Genbank no. BA000016.3) were used as templates for primer design and PCR amplification. The *rstA* ortholog from *C. acetobutylicum* was synthesized by Genscript (Piscataway, NJ). The *Streptococcus pyogenes* CRISPR-Cas9 system, which has been modified for use in *C. difficile* (31), was used to create a non-polar deletion of the *rstA* gene. The 630Δ*erm* and RT1075 (*sigD*::*erm*) strains containing the *rstA*-targeted CRISPR-Cas9 plasmid (MC1133 and MC1193, respectively) were grown overnight in TY medium with 5 μg/ml thiamphenicol. The next morning the cultures were backdiluted into fresh TY medium supplemented with 5 μg/ml thiamphenicol and 100 ng/ml anhydrous tetracycline for 24 h to induce expression of the CRISPR-Cas9 system. A small aliquot of this culture was streaked onto BHIS plates, and colonies were screened by PCR for the presence or absence of the *rstA* allele

### Mapping the *rstA* transcriptional start with 5’ rapid amplification of cDNA ends (5’ RACE)

DNase-I treated RNA from the *rstA*::*erm* mutant (MC391) was obtained as described above. 5’ RACE was performed using the 5’/3’ RACE Kit, Second Generation (Roche), following the manufacturer’s instructions as previously reported (32). Briefly, first strand cDNA synthesis was performed using the *rstA-*specific primer oMC982, followed by purification with the High Pure PCR Product Purification Kit (Roche). After subsequent poly(A)-tailing of first strand cDNA, PCR amplification was performed using an oligo(T) primer and the *rstA-*specific primer oMC983 with Phusion DNA Polymerase (NEB). The resulting PCR products were purified from a 0.7% agarose gel (Qiagen) and TA cloned into pCR2.1 (Invitrogen) using the manufacturer’s supplied protocols. Plasmids were isolated and sequenced (Eurofins MWG Operon) to determine the transcriptional start site (−32 bp from translational start site; n = 7).

### Sporulation assays

*C. difficile* cultures were grown in BHIS medium supplemented with 0.1% taurocholate and 0.2% fructose until mid-exponential phase (*i.e.*, an optical density at 600 nm (OD_600_) of 0.5), and 0.25 ml were spotted onto 70:30 sporulation agar as a lawn (27). After 24 h growth, ethanol resistance assays were performed as previously described (33, 34). Briefly, the cells were scraped from plates after 24 hours (H_24_) and suspended in BHIS medium to an OD_600_ of 1.0. The total number of vegetative cells per ml was determined by immediately serially diluting and applying the resuspended cells to BHIS plates. Simultaneously, a 0.5 ml aliquot was mixed with 0.3 ml 95% ethanol and 0.2 ml dH_2_O, to achieve a final concentration of 28.5% ethanol, vortexed and incubated for 15 min to eliminate all vegetative cells; ethanol-treated cells were subsequently serially diluted in 1X PBS + 0.1% taurocholate and applied to BHIS + 0.1% taurocholate plates to determine the total number of spores. After at least 36 h of growth, CFU were enumerated and the sporulation frequency was calculated as the total number of spores divided by the total number of viable cells (spores + vegetative cells). A *spo0A* mutant (MC310) was used as a negative sporulation control. Statistical significance was determined using a one-way ANOVA, followed by Dunnett’s multiple-comparison test (GraphPad Prism v6.0), to compare sporulation efficiency to the *rstA* mutant.

### Alkaline phosphatase activity assays

*C. difficile* strains containing the reporter fusions listed in **Table S1** were grown and harvested on either 70:30 sporulation agar at H_8_ or from TY liquid medium in stationary phase (T_3_ or H_24_). Alkaline phosphatase assays were performed as described previously (35) with the exception that no chloroform was used for cell lysis. Technical duplicates were averaged, and the results are presented as the mean and standard error of the mean of three biological replicates. The two-tailed Student’s *t* test was used to compare the activity in the *rstA* mutant to the parent strain.

### Biotin pulldown assays

Biotin pulldown assays were performed as described in Jutras *et. al*. 2012 (36). Briefly, an excess of biotin-labeled DNA bait was coupled to streptavidin-coated magnetic beads (Invitrogen) in B/W Buffer, and the bead-DNA complexes were washed with TE Buffer to remove unbound DNA. To prepare cell lysates, *C. difficile* expressing either RstA-FLAG (MC1004) or RstAΔHTH-FLAG (MC1028) in the *rstA* background were grown to mid-log phase (OD_600_ = 0.5) in 500 ml TY medium, pH 7.4, pelleted, rinsed with sterile water and stored at −80°C overnight. The pellets were suspended in 4.5 ml BS/THES Buffer and lysed by cycling between a dry ice/ethanol bath and a 37°C water bath. The cell lysates were vortexed for 1 min to shear genomic DNA, and cell debris was pelleted at 15K rpm for 15 min at 4°C. The supernatant, along with 10 μg/ml salmon sperm DNA as a nonspecific competitor, was then applied to the bead-DNA complexes and rotated for 30 min at room temperature. This incubation was repeated once with additional supernatant for two total incubations. The bead-DNA-protein complexes were washed extensively with BS/THES Buffer supplemented with, and then without, 10 μg/ml salmon sperm DNA to remove nonspecific proteins. The remaining bound protein was eluted with 250 mM NaCl in Tris-HCl, pH 7.4, and the eluates were immediately analyzed by SDS-PAGE and Western blot using FLAG M2 antibody (Sigma; see below). Each DNA bait fragment was tested in at least two independent experiments.

### Western blot analysis

The indicated *C. difficile* strains were grown in TY medium at 37°C and harvested at 24 h (34). Total protein was quantitated using the Pierce Micro BCA Protein Assay Kit (Thermo Scientific), and 8 μg of total protein was separated by electrophoresis on a precast 4-15% TGX gradient gel (Bio-Rad) and then transferred to a 0.45 μm nitrocellulose membrane. Western blot analysis was conducted with either mouse anti-TcdA (Novus Biologicals) or mouse anti-FLAG (Sigma) primary antibody, followed by goat anti-mouse Alexa Fluor 488 (Life Technologies) secondary antibody. Imaging and densitometry were performed with a ChemiDoc and Image Lab Software (Bio-Rad), and a one-way ANOVA, followed by Dunnett’s multiple-comparison test, was performed to assess statistical differences in TcdA protein levels between the *rstA* mutant and each *rstA* overexpression strain (GraphPad Prism v6.0). At least three biological replicates were analyzed for each strain, and a representative Western blot image is shown.

### Quantitative reverse transcription PCR analysis (qRT-PCR)

*C. difficile* was cultivated in TY medium and harvested at T_3_ (defined as three hours after the start of transition phase; OD_600_ = 1.0). Aliquots of 3 ml culture were immediately mixed with 3 ml ice-cold ethanol:acetone (1:1) and stored at −80°C. RNA was purified and DNase-I treated (Ambion) as previously described (37-39), and cDNA was synthesized using random hexamers (39). Quantitative reverse-transcription PCR (qRT-PCR) analysis, using 50 ng cDNA per reaction and the SensiFAST SYBR & Fluorescein Kit (Bioline), was performed in technical triplicates on a Roche Lightcycler 96. cDNA synthesis reactions containing no reverse transcriptase were included as a negative control to ensure no genomic DNA contamination was present. Results are presented as the mean and standard error of the means of three biological replicates. Statistical significance was determined using a one-way ANOVA, followed by Dunnett’s multiple-comparison test (GraphPad Prism v6.0), to compare transcript levels between the *rstA* mutant and each *rstA* overexpression strain.

## RESULTS

### RstA autoregulates its gene transcription via an inverted repeat overlapping the promoter

Our previous work provided preliminary genetic evidence that the N-terminal putative helix-turn-helix DNA-binding domain was necessary for regulation of toxin gene expression but was dispensable for sporulation initiation (14). However, further work with the recombinant His-tagged RstA proteins revealed that the constructs were expressed at low levels and were not detectable by Western blot in *C. difficile* lysates (data not shown). We created a new series of tagged proteins, possessing the 3XFLAG tag on the C-terminal end, and found that these were stably expressed and easily detected in *C. difficile rstA*::*erm* lysates (**Fig. S2A**). Corroborating our previous data, expression of the wild-type RstA, the full-length FLAG-tagged RstA and the truncated RstAΔHTH-FLAG-tagged allele complemented sporulation in the *rstA* mutant (**Fig. S2B**). As previously observed, only full-length RstA restored toxin production to wild-type levels in the *rstA* background (**Fig. S2C**), confirming that the helix-turn-helix motif within the DNA-binding domain is essential for RstA-dependent control of toxin production.

We hypothesized that RstA directly binds to DNA to control toxin gene expression and transcription of additional target genes. This interaction is predicted to occur via the putative DNA-binding domain, as observed for other RRNPP transcriptional regulators (40-42). To test this, we first defined the *rstA* promoter region and probed the DNA-binding capability of RstA within the promoter. The transcriptional start of *rstA* was identified at −32 bp upstream from the translational start using 5’ RACE. Corresponding σ^A^-10 and −35 consensus sequences were detected immediately upstream of this transcriptional start site (**Fig. 1A, B**). To verify the mapped promoter and to determine if any additional promoters are present that drive *rstA* transcription, a series of promoter fragments fused to the *phoZ* reporter gene was created, and alkaline phosphatase (AP) activity was measured in the 630Δ*erm* and *rstA*::*erm* mutant. As previously observed, the full-length 489 bp *rstA* promoter fragment exhibited a 1.8-fold increase in activity in the *rstA* mutant compared to the parent strain, indicating RstA-dependent repression (**Fig. 1C**, (14)). The truncated promoter fragments, P*rstA*_291_ and P*rstA*_231_, produced a similar fold change in activity in the *rstA* mutant and parent strain, as was observed for the full-length promoter. However, reporter activity was lower in the P*rstA*_115_ fragment compared to the longer fragments, suggesting that an enhancer sequence or an additional RstA-independent transcriptional activator is located between −231 bp to −115 bp upstream of the *rstA* open reading frame. A promoter fragment reporter fusion containing 380 bp of sequence upstream from the mapped *rstA* promoter (from −489 bp to −112 bp; IR; **Fig. 1A**) was inactive, indicating that an additional promoter is not located within this region. Altogether, the data demonstrate that the mapped σ^A^-dependent promoter drives *rstA* expression and that RstA can repress transcription from this promoter.

**Figure 1.**
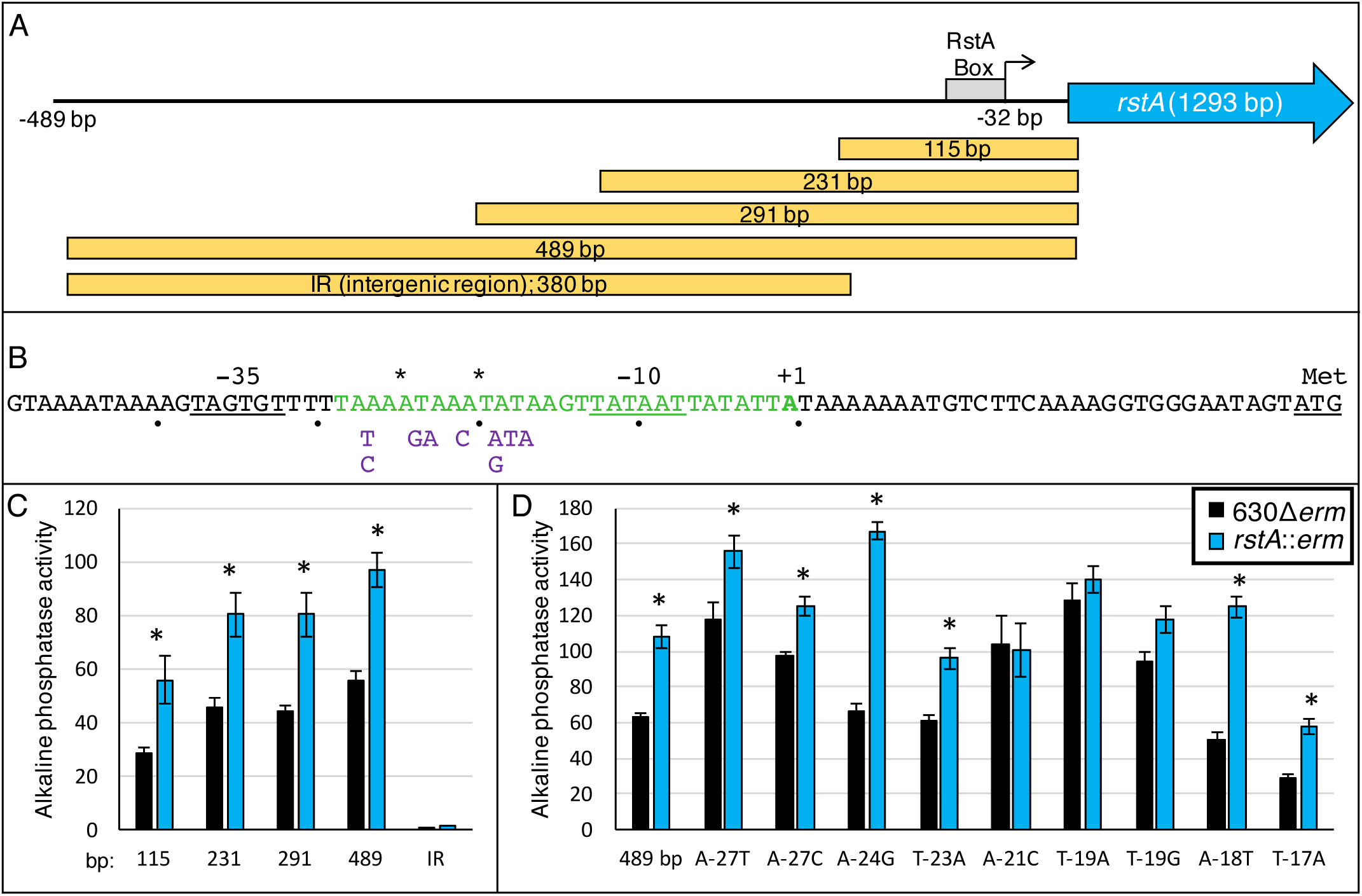
RstA controls its gene expression through an inverted repeat sequence overlapping the *rstA* promoter. (**A**) A schematic of the *rstA* promoter region denoting the general location of the putative RstA-box, the transcriptional start (32 bp upstream from the start codon; represented by the bent arrow) and the *rstA* open reading frame (not to scale). The yellow boxes indicate the location and size of promoter fragments constructed for the *phoZ* reporter fusions in panel C. (**B**) The *rstA* promoter, marked by +1, overlaps a 21 bp imperfect inverted repeat, colored green. The asterisks above the sequence mark the mismatched nucleotides within the inverted repeat. The −10 and −35 consensus sequences and the ATG start codon are underlined. The nucleotides below the sequence represent the substitutions tested in panel D. Alkaline phosphatase (AP) activity of the P*rstA*::*phoZ* reporter fusions of various lengths, including the upstream intergenic region (IR; −489 bp to −112 relative to the translational start) of *rstA* (**C**; P*rstA*_115_ (MC979/MC980); P*rstA*_231_ (MC1010/MC1011); P*rstA*_291_ (MC1012/MC1013); P*rstA*_489_ (MC773/MC774); P*rstA*_IR_ (MC1008/MC1009)) or of the full-length P*rstA*::*phoZ* promoter with various nucleotide substitutions (**D**; P*rstA*_489_ (MC773/MC774); P*rstA*_A-27T_ (MC830/MC831); P*rstA*_A-27C_ (MC856/MC857); P*rstA*_A-24G_ (MC858/MC859); P*rstA*_T-23A_ (MC832/MC833); P*rstA*_A-21C_ (MC860/MC861); P*rstA*_T-19A_ (MC834/MC835); P*rstA*_T-19G_ (MC862/MC863); P*rstA*_A-18T_ (MC836/MC837); P*rstA*_T-17A_ (MC838/MC839)) in 630Δ*erm* and the *rstA*::*erm* mutant (MC391), respectively, grown on 70:30 sporulation agar at H_8_. The means and standard error of the means of four biological replicates are shown. *, *P* < 0.05, using Student’s *t-*test compared to the activity observed in the 630Δ*erm* parent strain for each promoter construct.

The results obtained from the promoter-reporter fusions suggested that RstA binding was likely to occur within the 115 bp upstream of the translational start site. A 29 bp imperfect inverted repeat was identified within the predicted P*rstA* −10 consensus sequence, suggesting a possible regulatory binding site within this region (**Fig. 1B**). To determine whether this sequence serves as an RstA recognition site, we created a series of single nucleotide substitutions within the inverted repeat in the 489 bp P*rstA* reporter fusion, avoiding conserved residues required for RNAP-holoenzyme recognition (43). Most of the single nucleotide substitutions did not significantly alter reporter activity compared to the wild-type P*rstA* reporter (**Fig. 1D**). However, three nucleotide substitutions in two positions, A-21 and T-19, abolished RstA repression in the parent strain, increasing reporter activity to match that of the *rstA*::*erm* mutant. These data suggest that the A-21 and T-19 nucleotides are important for RstA binding to the *rstA* promoter.

### RstA inhibits toxin and motility gene transcription

Regulatory control of toxin gene expression in *C. difficile* involves multiple sigma factors and transcriptional regulators, which ensure toxin production occurs in the appropriate environmental conditions (13). Our previous work (14) demonstrated that an *rstA*::*erm* mutant has increased transcription of the *C. difficile* toxin genes, *tcdA* and *tcdB*, the toxin-specific sigma factor, *tcdR*, and the flagellar-specific sigma factor, *sigD*, which is required for motility and directs *tcdR* transcription (11, 12). To determine whether RstA is involved directly in repressing transcription of these genes, we first constructed *phoZ* reporter fusions with the promoter regions for each gene and examined RstA-dependent transcriptional activity.

The *tcdR* promoter region contains four identified independent promoter elements: a σ^A^- dependent promoter (−16 bp from the translational start), a σ^D^-dependent promoter (−76 bp from the translational start) and two putative σ^TcdR^ promoters farther upstream (**Fig. 2A**; (11, 12, 44- 46)). Expression of the *tcdR* gene is relatively low in *C. difficile* (11, 45, 47), at least in part due to repression by CodY and CcpA binding throughout the *tcdR* promoter region under nutrient-rich conditions (8, 9, 37, 46, 48). We examined each of the promoter elements within P*tcdR* to determine if RstA affects transcription from these promoters. A series of reporter fusions was created for each of the promoter elements, which were examined in the *rstA*::*erm* mutant and parent strain, respectively, and activity was measured after 24 h of growth in TY medium (**Fig. 2A**). A full-length 517 bp P*tcdR*::*phoZ* reporter and the two σ^TcdR^-dependent promoter fusions exhibited similar low reporter activities in the parent and the *rstA* strains (**Fig. 2B**). But, increased reporter activity was observed in the *rstA* mutant for the individual σ^A^-dependent and σ^D^-dependent promoter fusions. These results indicate that RstA impacts the function of these promoter elements and contributes to repression of *tcdR* transcription.

**Figure 2.**
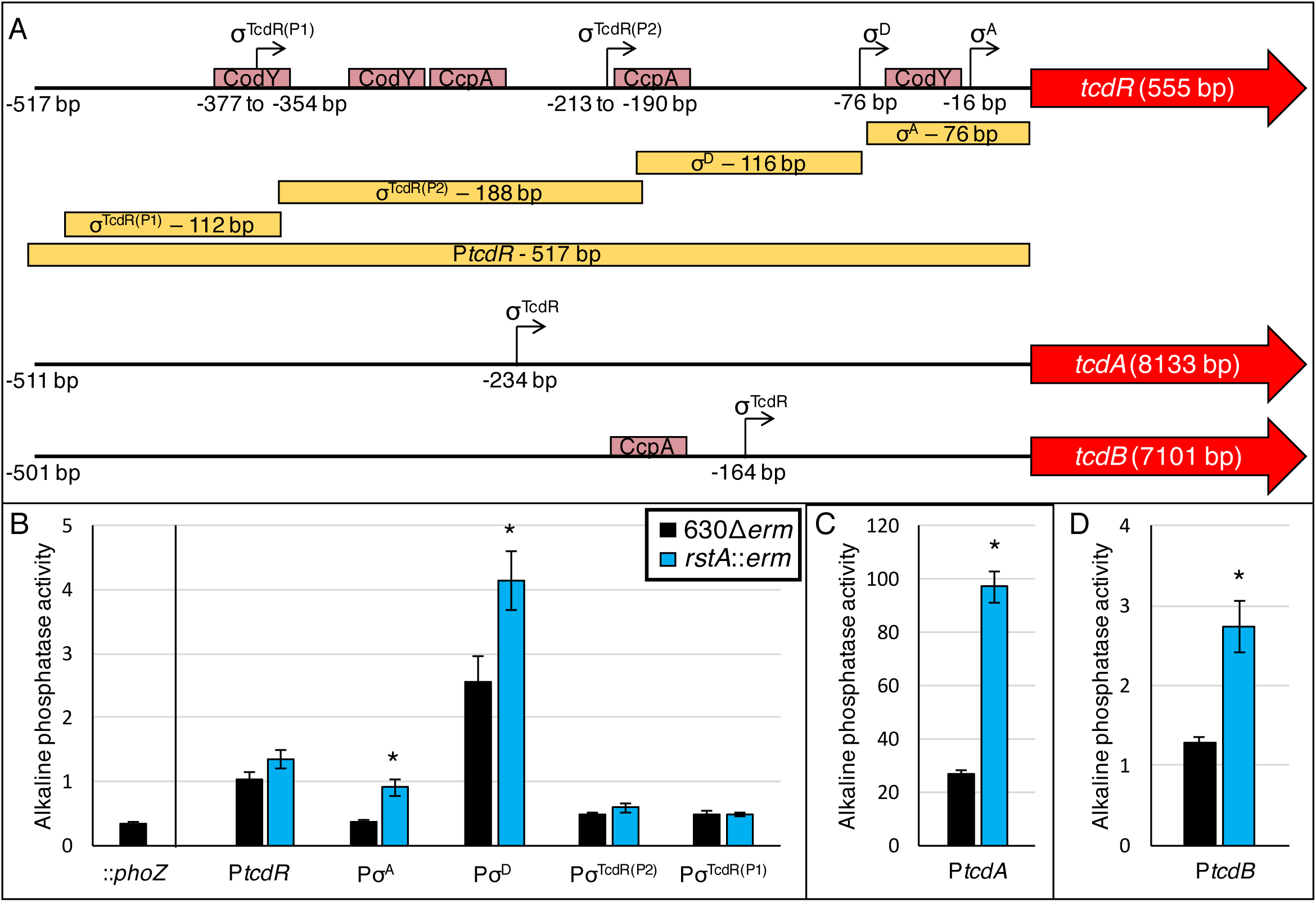
RstA inhibits toxin gene expression. (**A**) A schematic of the promoter regions of *tcdR, tcdA* and *tcdB* denoting the relative locations of the transcriptional start sites experimentally demonstrated (12, 45-47) and the open reading frame of all three genes (not to scale). Pale red boxes approximate CodY and CcpA binding sites within the toxin gene promoters (8, 9, 48). The yellow boxes indicate the location and size of the promoter fragments constructed for the *phoZ* reporter fusions in panels B-D. Alkaline phosphatase (AP) activity of (**B**) the P*tcdR*::*phoZ* reporter fusions of various lengths (promoterless *phoZ* (MC448); P*tcdR*_σA_ (MC1285/MC1286); P*tcdR*_σD_ (MC1145/MC1146); P*tcdR*_σTcdR(P2)_ (MC1147/MC1148); P*tcdR*_σTcdR(P1)_ (MC1149/MC1150)) and (**C**) the P*tcdA*::*phoZ* (−511 bp to −1 bp upstream of transcriptional start; MC1249/MC1250) or (**D**) P*tcdB*::*phoZ* (−531 bp to −31 bp upstream of transcriptional start; MC1251/MC1252)) reporter fusions in 630Δ*erm* and the *rstA*::*erm* mutant (MC391) grown in TY medium, pH 7.4 at H_24_. The means and standard error of the means of four biological replicates are shown. *, *P* < 0.05, using Student’s *t-*test compared to the activity observed in the 630Δ*erm* parent strain for each promoter construct.

We also examined RstA-dependent regulation of *tcdA* and *tcdB* transcription, both of which are expressed solely from σ^TcdR^-dependent promoters (**Fig. 2A;** (47, 49, 50)). P*tcdA* reporter activity was increased 3.6-fold and P*tcdB* activity was 2.1-fold greater in the *rstA* strain compared to the parent (**Fig. 2C, D**). Altogether, these data indicate that RstA represses toxin gene transcription at the individual gene level and through repression of *tcdR*.

SigD, also known as FliA or σ^28^, is a sigma factor that coordinates flagellar gene expression and directly activates *tcdR* gene expression (45). The *sigD* gene is located in a large, early-stage flagellar operon, that is transcribed from a σ^A^-dependent promoter located 496 bp upstream from the first gene of the operon, *flgB* (51). Interestingly, the *flgB* promoter sequences from two different *C. difficile* strains, the historical epidemic strain, 630, and a current epidemic strain, R20291, are identical from the σ^A^ promoter sequence through the translational start site, but diverge considerably upstream of this region (**Fig. S3**). No additional promoter elements were identified in the 630 or R20291 sequences upstream of the σ^A^-dependent promoter (**Fig. 3A**). To determine whether RstA influences *sigD* transcription through repression of P*flgB*, promoter reporter fusions representing each strain were constructed. As anticipated, activity of the 630Δ*erm* and R20291 P*flgB* reporters were higher in the *rstA* mutant compared to the parent strain (1.7-fold and 1.5-fold, respectively; **Fig. 3B**), indicating that RstA represses *flgB*, and consequently, *sigD* transcription.

**Figure 3.**
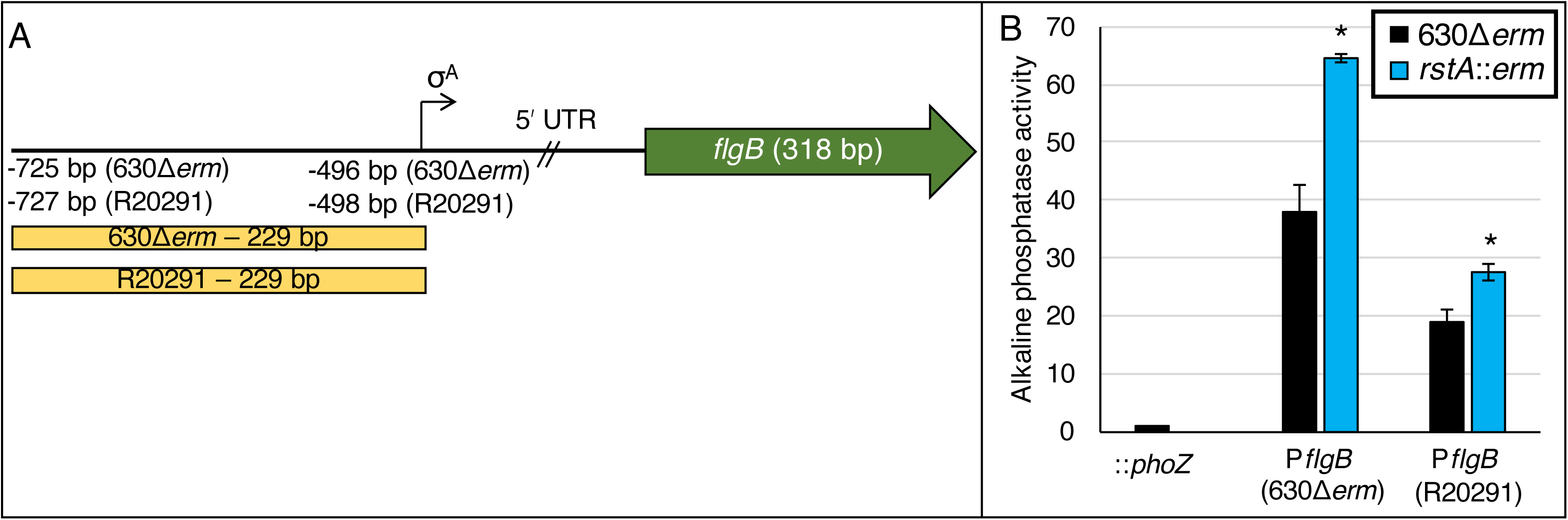
RstA represses expression of *flgB* reporter fusions. (**A**) A schematic of the *flgB* promoter regions for *C. difficile* 630 and R20291 strains. The transcriptional start site for the σ^A^- dependent promoter for 630 lies −496 bp upstream from the *flgB* translational start while the R20291 initiates transcription −498 bp upstream (51, 65). (**B**) Alkaline phosphatase (AP) activity of the promoterless::*phoZ* vector in 630Δ*erm* (MC1106) and P*flgB*_630Δ*erm*_::*phoZ* (MC1294/MC1295) and P*flgB*_R20291_::*phoZ* (MC1296/MC1297) reporter fusions in 630Δ*erm* and the *rstA*::*erm* mutant (MC391) grown in TY medium, pH 7.4 at T_3_ (three hours after the start of transition phase; OD_600_ = 1.0). The means and standard error of the means of three biological replicates are shown. *, *P* < 0.05, using Student’s *t-*test compared to the activity observed in the 630Δ*erm* parent strain for each promoter construct.

### RstA directly binds the *rstA, tcdR, flgB, tcdA* and *tcdB* promoters via the conserved helix-turn-helix DNA-binding domain

To determine whether RstA directly binds target DNA, a variety of *in vitro* electrophoretic gel shift assays were attempted, but no binding was observed in any condition tested. We considered that the lack of RstA-DNA interaction by gel shift may occur because of the absence of a cofactor, such as a quorum-sensing peptide, or because of a transient complex or oligomerization state. To overcome this obstacle, we performed biotin-labelled DNA pulldown assays to assess the DNA-binding capacity of RstA under native conditions. Biotinylated DNA was coupled to streptavidin beads as bait and incubated with cell lysates expressing either full-length RstA-FLAG or RstAΔHTH-FLAG protein. Specifically-bound proteins were eluted and analyzed by western blot using FLAG M2 antibody.

We first tested the ability of RstA to directly interact with its own promoter. RstA-FLAG protein was recovered using the wild-type *rstA* promoter region as bait, demonstrating specific interaction of the RstA protein (**Fig. 4A**). However, the P*rstA* fragment did not capture RstAΔHTH-FLAG protein, indicating that the conserved HTH domain of RstA is essential for DNA binding. In addition to the wild-type *rstA* promoter, the P*rstA* T-19A and P*rstA* A-21C variants that eliminated RstA-dependent regulation *in vivo* were used as bait (**Fig. 1D**). Both the P*rstA* T-19A and P*rstA* A-21C variants captured less RstA-FLAG than the wild-type promoter, suggesting that these nucleotides facilitate RstA interaction (**Fig. 4A**). Finally, the intergenic region upstream of the *rstA* promoter (**Fig. 1A; IR**) did not recover the full-length RstA-FLAG, indicating that RstA recognizes a specific DNA sequence within the promoter region. Altogether, these data demonstrate that RstA functions as a DNA-binding protein that directly and specifically binds its own promoter to repress transcription.

**Figure 4.**
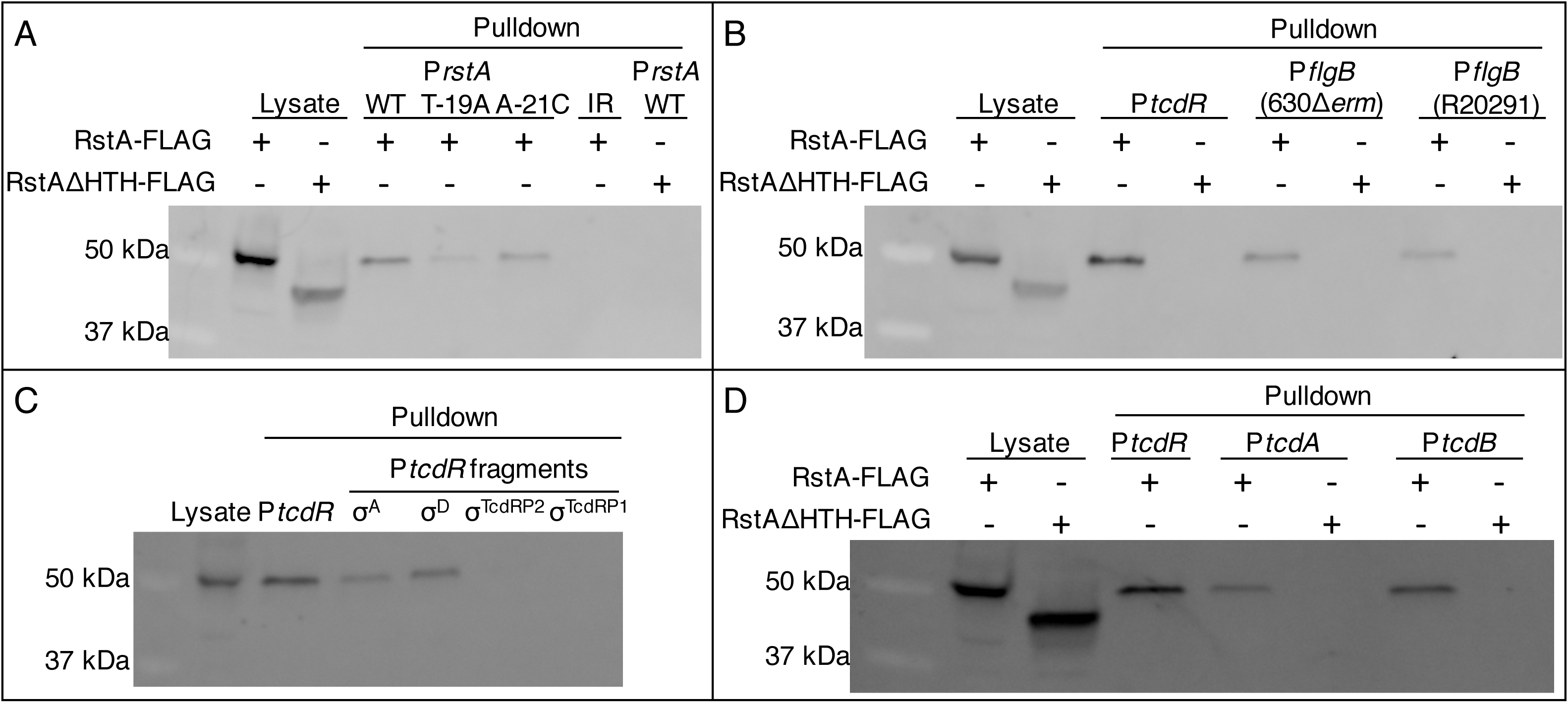
RstA binds to the *rstA, tcdR, flgB, tcdA* and *tcdB* promoters. Western blot analysis using FLAG M2 antibody to detect recombinant RstA-3XFLAG or RstAΔHTH-3XFLAG in cell lysates or following biotin-labeled DNA pull-down assays. As a control, cell lysate expressing the RstA-3XFLAG construct (MC1004) or the RstAΔHTH-3XFLAG construct (MC1028) is included in the first lane or two of each western blot shown. The biotin-labeled fragments used as bait are of (**A**) the 115 bp wild-type, T-19A or A-21C *rstA* promoters or of the 380 bp intergenic region upstream of the *rstA* promoter (IR; see Fig. 2); (**B**) the full-length *tcdR* (446 bp) or the 630Δ*erm* or R20291 *flgB* (229 bp) promoters; (**C**) the full-length *tcdR* (446 bp), σ^A^-dependent (92 bp), σ^D^-dependent (116 bp), σ^TcdRP2^-dependent (188 bp), or σ^TcdRP1^-dependent (112 bp) promoters; or (**D**) the full-length *tcdR* (446 bp), *tcdA* (511 bp) or *tcdB* (501 bp) promoters. All promoter fragments were bound to streptavidin-coated magnetic beads and incubated with *C. difficile* cell lysates grown in TY medium to mid-log phase (OD_600_ = 0.5), expressing either the RstA-3XFLAG construct (MC1004) or the RstAΔHTH-3XFLAG construct (MC1028).

To determine if RstA directly binds DNA to repress the transcription of genes encoding toxin regulators, we examined RstA binding to the *flgB* and *tcdR* promoter regions. RstA-FLAG protein bound specifically to the full-length *tcdR* promoter region, as well as the 630 and R20291 *flgB* promoters (**Fig. 4B**). Again, the HTH domain was required for these RstA-promoter interactions. To identify which internal promoter elements directly interact with RstA, previously characterized *tcdR* promoter fragments were used as bait (**Fig. 2B**), with the exception of a longer σ^A^-dependent promoter fragment (92 bp rather than 76 bp) to limit potential steric hindrance of RstA binding due to the 5’ biotin label. This longer 92 bp P*tcdR*(σA) fragment exhibited the same RstA-dependent regulation in reporter assays as the 76 bp reporter (**Fig. S4**). RstA-FLAG bound to the σ^A^-dependent and σ^D^-dependent *tcdR* promoter fragments but was not recovered from either of the σ^TcdR^-dependent promoters (**Fig. 4C**), corroborating the reporter fusion results that demonstrated RstA repression of only the σ^A^-dependent and σ^D^- dependent *tcdR* promoter elements.

DNA pulldown assays were also performed to ascertain if RstA directly binds to the *tcdA* and *tcdB* promoters. Both of the toxin promoters captured the full-length RstA-FLAG protein and failed to recover the RstAΔHTH-FLAG protein (**Fig. 4D**). These data provide direct biochemical evidence that RstA represses *tcdR, tcdA* and *tcdB* transcription by binding to the promoter regions of these genes.

### RstA represses toxin gene expression independently of SigD-mediated toxin regulation

Our data indicate that RstA represses toxin gene expression directly by binding to the *tcdA* and *tcdB* promoter regions, and indirectly by repressing transcription of the sigma factors *tcdR* and *sigD*, which activate toxin gene expression. To confirm genetically that RstA represses toxin gene expression independently of *sigD*, we created an *rstA sigD* double mutant and examined the impact of each mutation on toxin production. To aid in construction of a *rstA sigD* double mutant, we utilized the recently developed CRISPR-Cas9 system modified for use in *C. difficile* to create an unmarked, non-polar deletion of *rstA* in the 630Δ*erm* and *sigD*::*erm* backgrounds (**Fig. S5**) (31). TcdA protein levels were ~3-fold higher in the *rstA sigD* double mutant compared to the *sigD* mutant (**Fig. 5A**), indicating that RstA represses toxin production independently of SigD. Overexpression of *rstA* in the *rstA sigD* mutant returned TcdA protein to the levels found in the *sigD* mutant. Likewise, *sigD* overexpression in the *rstA sigD* mutant restored TcdA to wild-type levels, further supporting that SigD and RstA regulate toxin production independently (**Fig. 5A**). In addition, transcript levels of *tcdA, tcdB,* and *tcdR* were increased in the *rstA sigD* mutant compared to the *sigD* mutant (**Fig. 5B**), mirroring the TcdA protein results. Altogether, these data provide further evidence that RstA is major regulator of toxin production that directly and indirectly represses toxin gene expression independently of SigD.

**Figure 5.**
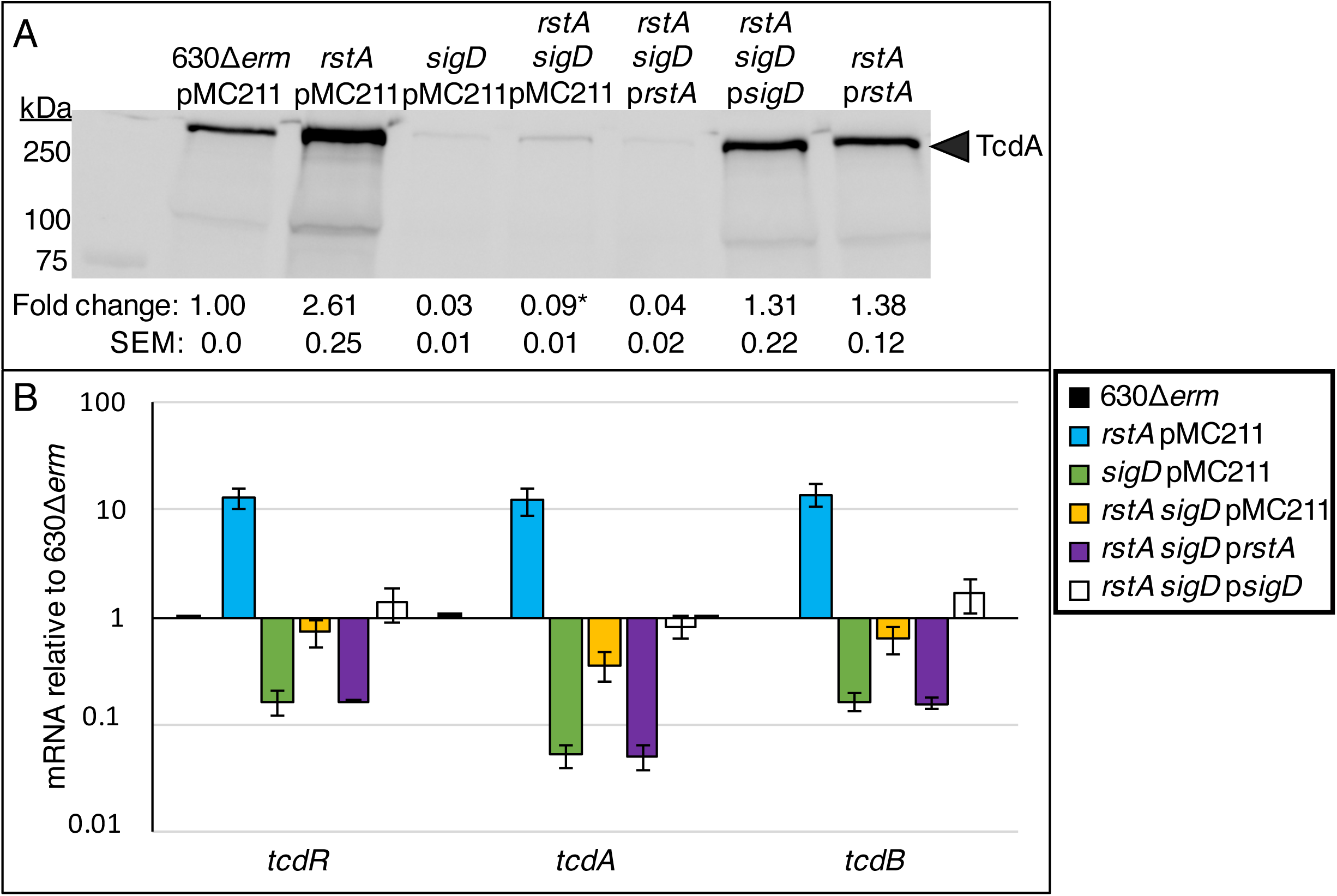
RstA represses toxin gene expression independently of SigD-mediated regulation. (**A**) Western blot analysis of TcdA in 630Δ*erm* pMC211 (MC282; vector control), *rstA* pMC211 (MC1224; vector control), *sigD*::*erm* pMC211 (MC506; vector control), *rstA sigD*::*erm* pMC211 (MC1281), *rstA sigD*::*erm* p*PcprA-rstA* (MC1282), *rstA sigD*::*erm* p*PcprA-sigD* (MC1283) and *rstA* pP*cprA-rstA* (MC1225) grown in TY medium supplemented with 2 μg/ml thiamphenicol and 1 μg/ml nisin, pH 7.4, at 24 h. (**B**) qRT-PCR analysis of *tcdR, tcdA*, and *tcdB* transcript levels in 630Δ*erm* pMC211 (MC282; vector control), *rstA* pMC211 (MC1224; vector control), *sigD*::*erm* pMC211 (MC506; vector control), *rstA sigD*::*erm* pMC211 (MC1281), *rstA sigD*::*erm* p*PcprA-rstA* (MC1282) and *rstA sigD*::*erm* p*PcprA-sigD* (MC1283) grown in TY medium supplemented with 2 μg/ml thiamphenicol and 1 μg/ml nisin, pH 7.4, at T_3_ (three hours after the entry into stationary phase). The means and standard error of the means of three biological replicates are shown. *, *P* < 0.05, Student’s *t*-test between *sigD::erm* pMC211 and *rstA sigD*::*erm* pMC211.

### RstA DNA-binding activity requires a species-specific predicted quorum-sensing domain

The observation that RstA does not bind to target DNA in the tested *in vitro* conditions, but does bind DNA in cell lysates, suggests that a co-factor is required for RstA DNA-binding activity. We hypothesize that a small, quorum-sensing peptide serves as an activator for RstA DNA-binding, as has been observed for other members of the RRNPP family (23-25, 52-54). To test this, we expressed RstA orthologs of other clostridial species (**Fig. S6**), including *Clostridium acetobutylicum* (*C.a.*), *Clostridium perfringens* (*C.p.*), and *Clostridium* (*Paeniclostridium*) *sordellii* (*C.s.*) in the *C. difficile rstA* mutant background. Only the *C.p.* RstA was stably produced in *C. difficile* (**Fig. S7**). However, expression of the *C. perfringens rstA* ortholog failed to restore TcdA protein to wild-type levels (**Fig. 6A**). *C.p.* RstA may be unable to repress *C. difficile* toxin production because the *C.p.* DNA-binding domain cannot recognize the *C. difficile* DNA target sequences and/or because the DNA-binding activity of *C.p.* RstA is not functional in *C. difficile*. To distinguish between these possibilities, we constructed a chimeric protein containing the *C. perfringens* DNA-binding domain (M1-Y51) fused to the C-terminal domains of the *C. difficile* RstA protein (herein known as CpHTH-CdCterminalFLAG) and examined the function of this chimeric RstA in the *C. difficile rstA* mutant. The RstA chimera restored *C. difficile* TcdA levels to those observed in the parent strain (**Fig. 6A**), indicating that the *C. perfringens* DNA-binding domain is functional in *C. difficile*. These data strongly suggest that the C-terminal portion of RstA responds to species-specific intracellular signals to control the N-terminal DNA-binding activity.

Finally, we assessed the ability of a *C. perfringens* RstA to complement the low sporulation frequency of the *C. difficile rstA* mutant. Overexpressing the full-length *C. perfringens* RstA did not complement sporulation in the *C. difficile rstA* mutant (**Fig. 6B**). Unexpectedly, a hypersporulation phenotype was observed when the CpHTH-CdC-terminalFLAG RstA chimera was expressed in the *rstA* mutant (**Fig. 6B**), indicating that the chimeric *C.p.-C.d.* RstA promotes *C. difficile* sporulation to even higher levels than the native *C. difficile* RstA. This hypersporulation phenotype suggests that the *C.p.* HTH portion of the chimeric RstA protein alters the structure or activity of RstA to increase the positive effect on early sporulation events. These data warrant further investigation into the molecular mechanisms by which the C-terminal domains of RstA cooperate with the DNA-binding domain to promote sporulation.

**Figure 6.**
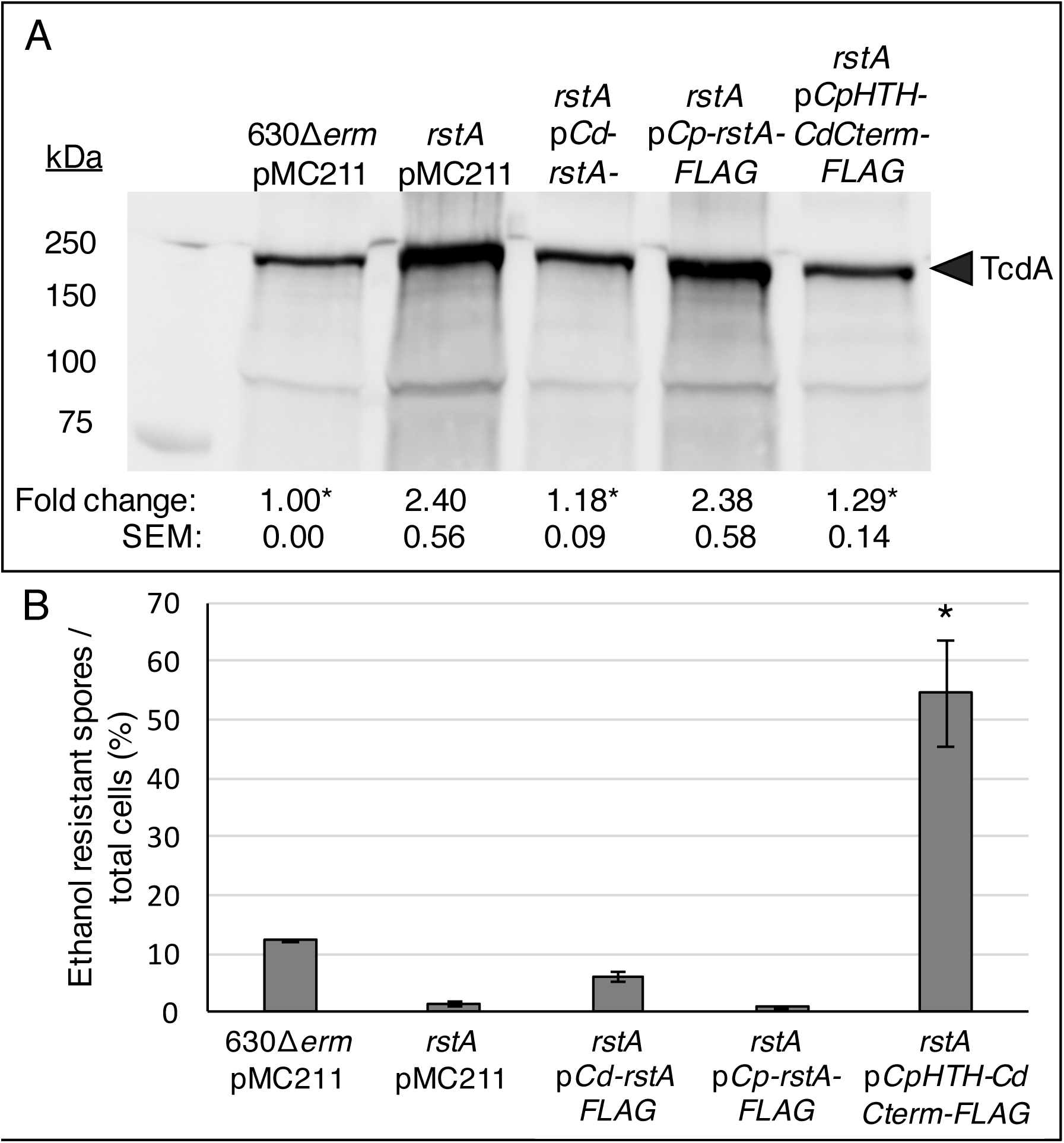
A hybrid *rstA* construct containing the *C. perfringens* DNA-binding domain with the *C. difficile* quorum-sensing-like domain complements *C. difficile rstA* toxin production and sporulation. (**A**) Western blot analysis of TcdA in 630Δ*erm* pMC211 (MC282; vector control), *rstA*::*erm* pMC211 (MC505; vector control), *rstA*::*erm* pP*cprA-rstA3XFLAG* (MC1004), *rstA*::*erm* pP*cprA-Cp-rstA3XFLAG* (MC1324) and *rstA*::*erm* pP*cprA-CpHTHCdCterminal3XFLAG* (MC1257) grown in TY medium, pH 7.4, at H_24_. (**B**) Ethanol resistant spore formation of 630Δ*erm* pMC211 (MC282; vector control), *rstA*::*erm* pMC211 (MC505; vector control), *rstA*::*erm* pP*cprA-rstA3XFLAG* (MC1004), *rstA*::*erm* pP*cprA-Cp-rstA3XFLAG* (MC1324) and *rstA*::*erm* pP*cprA-CpHTHCdCterminal3XFLAG* (MC1257) grown on 70:30 sporulation agar supplemented with 2 μg/ml thiamphenicol and 1 μg/ml nisin. Sporulation frequency is calculated as the number of ethanol-resistant spores divided by the total number of cells enumerated at H_24_ as detailed in the Methods and Materials. The means and standard error of the means of at least three independent biological replicates are shown; asterisks represent *P* ≤ 0.05 by one-way ANOVA followed by Tukey’s multiple comparison’s test compared to *rstA* pMC211 (MC505).

## DISCUSSION

The production of exotoxins and the ability to form quiescent endospores are two essential features of *C. difficile* pathogenesis. The regulatory links between toxin production and spore formation are complex and poorly understood. Some conserved sporulation regulatory factors, including Spo0A, CodY, and CcpA, strongly influence toxin production, yet some of these regulatory effects appear to be strain-dependent or are indirect (8, 48, 55-57). Further, additional environmental conditions and metabolic signals, such as temperature and proline, glycine and cysteine availability (5, 6, 10, 58), impact toxin production independently of these regulators, revealing the possibility that additional unknown factors are directly involved in toxin regulation (13). The recently discovered RRNPP regulator, RstA, represses toxin production and promotes spore formation, potentially providing a direct and inverse link between *C. difficile* spore formation and toxin biogenesis (14).

In this study, we show that RstA is a major, direct transcriptional regulator of *C. difficile* toxin gene expression. RstA inhibits toxin production by directly binding to the *tcdA* and *tcdB* promoters and repressing their transcription. RstA reinforces this repression by directly downregulating gene expression of *tcdR*, which encodes the sole sigma factor that drives *tcdA* and *tcdB* transcription. Finally, RstA directly represses the *flgB* promoter, inhibiting gene expression of the flagellar-specific sigma factor, SigD. SigD activates motility gene transcription but is also required for full expression of *tcdR* (11, 12). RstA repression of each major component in the toxin regulatory pathway creates a multi-tiered network in which RstA directly and indirectly controls *tcdA* and *tcdB* gene expression (**Fig. 7**).

**Figure 7.**
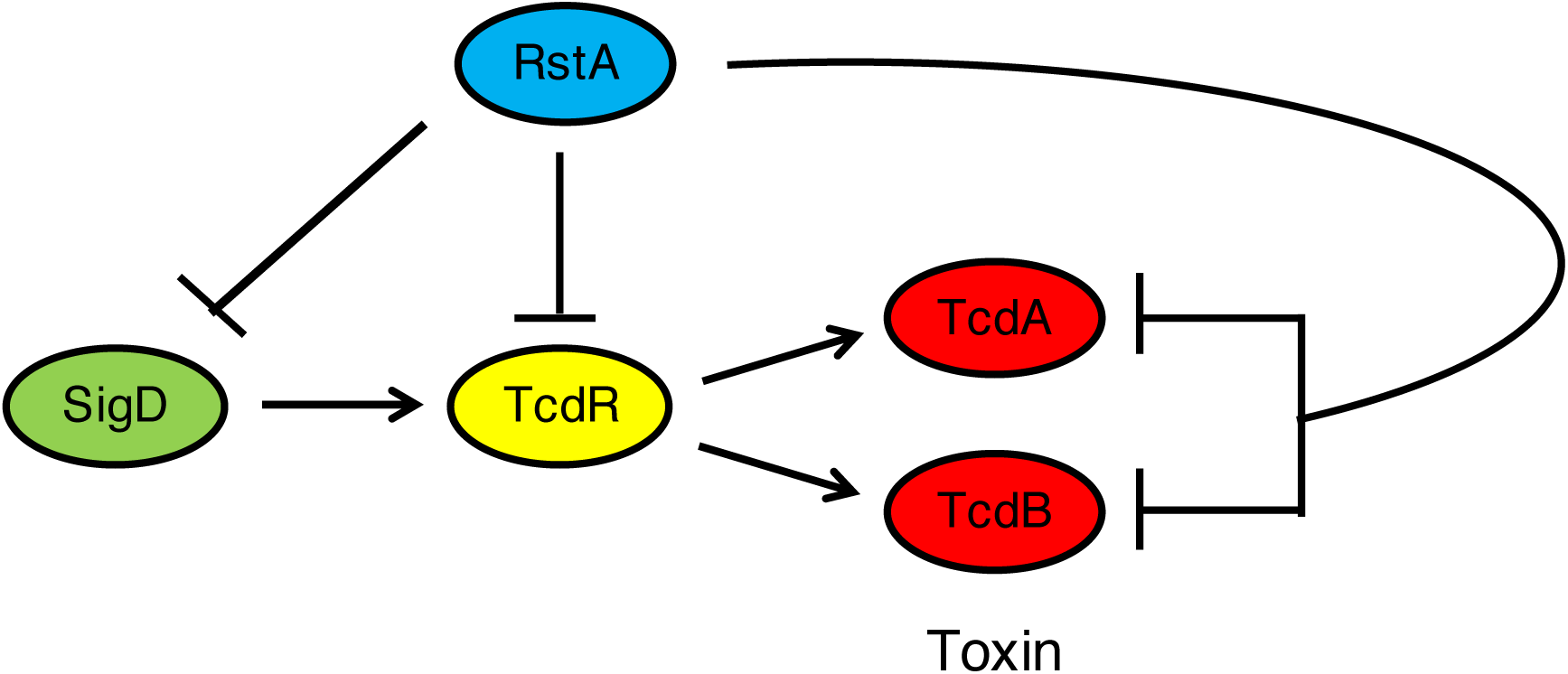
Model of RstA-mediated repression of *C. difficile* toxin production. SigD, the flagellar-specific sigma factor, directly induces gene transcription of *tcdR*, the toxin-specific sigma factor. Toxin gene expression is then directed by TcdR. RstA inhibits production of TcdA and TcdB by directly binding to and repressing transcription of *sigD, tcdR, tcdA* and *tcdB*, creating a complex, multi-tiered regulatory network to ensure that the toxin gene expression is appropriately timed in response to the signal(s) that activate RstA.

RstA is the third characterized transcriptional repressor that directly binds to promoter regions for *tcdR, tcdA,* and *tcdB*. The two others, CodY and CcpA, exhibit ~10-fold higher affinity *in vitro* for the *tcdR* promoter than for the *tcdA* and *tcdB* promoters (8, 9, 48), suggesting that CodY and CcpA repress toxin gene expression primarily through repression of *tcdR* transcription. Similarly, RstA appears to bind with greater affinity to the *tcdR* promoter than to the *tcdA* and *tcdB* promoters, although we caution that biotin pulldown results are semi-quantitative. The *in* vivo contribution of this reinforced repression of *tcdA* and *tcdB* transcription by CodY, CcpA, and RstA remains unknown. Interestingly, recent evidence has demonstrated that *tcdR* gene expression serves as a bistable switch that determines whether individual *C. difficile* cells within a population produce TcdA and TcdB, creating a divided population of toxin-OFF and toxin-ON cells (59). TcdR governs this bistability state by maintaining low basal expression levels, allowing for small changes to result in stochastic gene expression, and by positively regulating its own expression, establishing a positive feedback loop that bolsters the toxin-ON state (59). CodY was found to influence the population so that fewer cells produced toxin, but CcpA and RstA were not tested (59). We predict that both CcpA and RstA would bias the population of cells to a toxin-OFF state. Altogether, the tight control of *tcdR* transcription, reinforced by direct repression of *tcdA* and *tcdB* transcription by CcpA, CodY, and RstA, results in the convergence of multiple regulatory pathways at the bistable *tcdR* promoter to coordinate toxin production in response to nutritional and species-specific signals. This complex regulation ensures that the energy-intensive process of toxin production is initiated only to benefit the bacterium under the appropriate conditions.

Importantly, RstA is the first transcriptional regulator demonstrated to directly control *flgB* transcription initiation. To date, none of previously identified regulators of *flgB* expression, including Spo0A, SigH, Agr, Hfq, SinR and SinR’, have been shown to bind promoter DNA and regulate flagellar gene expression through transcription initiation (55, 60-63). *flgB* expression is further regulated post-transcriptionally via a c-di-GMP riboswitch or flagellar switch, both of which are located within the large, 496 bp 5′ untranslated region (51, 64, 65); however, the impact of RstA-mediated repression of *flgB* gene expression through additional pathways has not yet been explored.

Although we have identified several direct RstA targets, the sequence required to recruit RstA to target promoters remains unclear. The *rstA* promoter contains a near-perfect inverted repeat; however, this sequence is AT-rich, as is the case for many *C. difficile* promoters. Imperfect inverted repeats were also found overlapping the −35 consensus sequences of the *tcdA, tcdB, flgB* and σ^A^-dependent *tcdR* promoters, and immediately upstream of the σ^D^- dependent *tcdR* promoter (**Fig. S8**), suggesting that RstA inhibits transcription at these promoters by sterically obstructing RNA polymerase docking. No clear consensus sequence defining an RstA-box is delineated from these sequences. Exhaustive attempts at ChIP-seq analysis to identify the *C. difficile* RstA regulon proved unsuccessful; however, our data imply that RstA is a transcriptional repressor that directly controls multiple *C. difficile* phenotypes, and additional targets within in the *C. difficile* genome seem likely.

The inability to recapitulate RstA-DNA-binding with purified RstA *in vitro* together with the functional analysis of *C. difficile* and *C. perfringens* full-length and chimeric proteins suggest that i) RstA DNA-binding activity requires a cofactor and ii) this cofactor is species-specific. Most RRNPP members are cotranscribed with their cognate quorum-sensing peptide precursor (19), but there are notable exceptions, including those encoded by unlinked genes (52, 66) and orphan receptors whose cognate ligands have not yet been discovered (67-69). Importantly, no type of ligand other than small, quorum-sensing peptides has been identified for RRNPP proteins. In addition to RstA, other quorum-sensing factors have been implicated in *C. Difficile* toxin production. The incomplete Agr1 and conserved Agr2 quorum-sensing systems induce toxin production through the production of a cyclic auto-inducer peptide (AIP) that is sensed extracellularly (61, 70, 71); however, it is highly unlikely that the extracellular AIP molecule directly interacts with the cytosolic RstA protein. In addition, the interspecies LuxS-derived autoinducer-2 (AI-2) quorum-sensing molecule was found to increase *C. difficile tcdA* and *tcdB* gene expression, but not *tcdR* (72), indicating that AI-2 does not signal through RstA either. Although there are no open reading frames adjacent to RstA that encode an apparent quorum-sensing peptide precursor, taken altogether, it is reasonable to presume that an unidentified quorum-sensing mechanism controls RstA-dependent regulation of *C. difficile* TcdA and TcdB production.

Finally, as RstA is necessary for efficient *C. difficile* spore formation, the possibility remains that species-specific signaling is required for RstA-dependent control of early sporulation and that RstA coordinates *C. difficile* toxin production and spore formation in response to the same signal(s). Elucidating the molecular mechanisms that govern RstA activity will provide important insights into the regulatory control between sporulation and toxin production, reveal host cues and conditions that lead to increased toxin production, and help delineate the early sporulation events that control *C. difficile* Spo0A phosphorylation and activation.

## ACKNOWLEDGEMENTS

We are grateful for the gift of *C. perfringens* S13 genomic DNA from Dr. Jorge Vidal. We give special thanks to Dr. Charles Moran and the members of McBride lab for helpful suggestions and discussions during the course of this work. This research was supported by the U.S. National Institutes of Health through research grants AI116933 and AI121684 to S.M.M, AI107029 to R.T., and AI120613 to B.R.A-F. The content of this manuscript is solely the responsibility of the authors and does not necessarily reflect the official views of the National Institutes of Health.

---

**Table S1.**
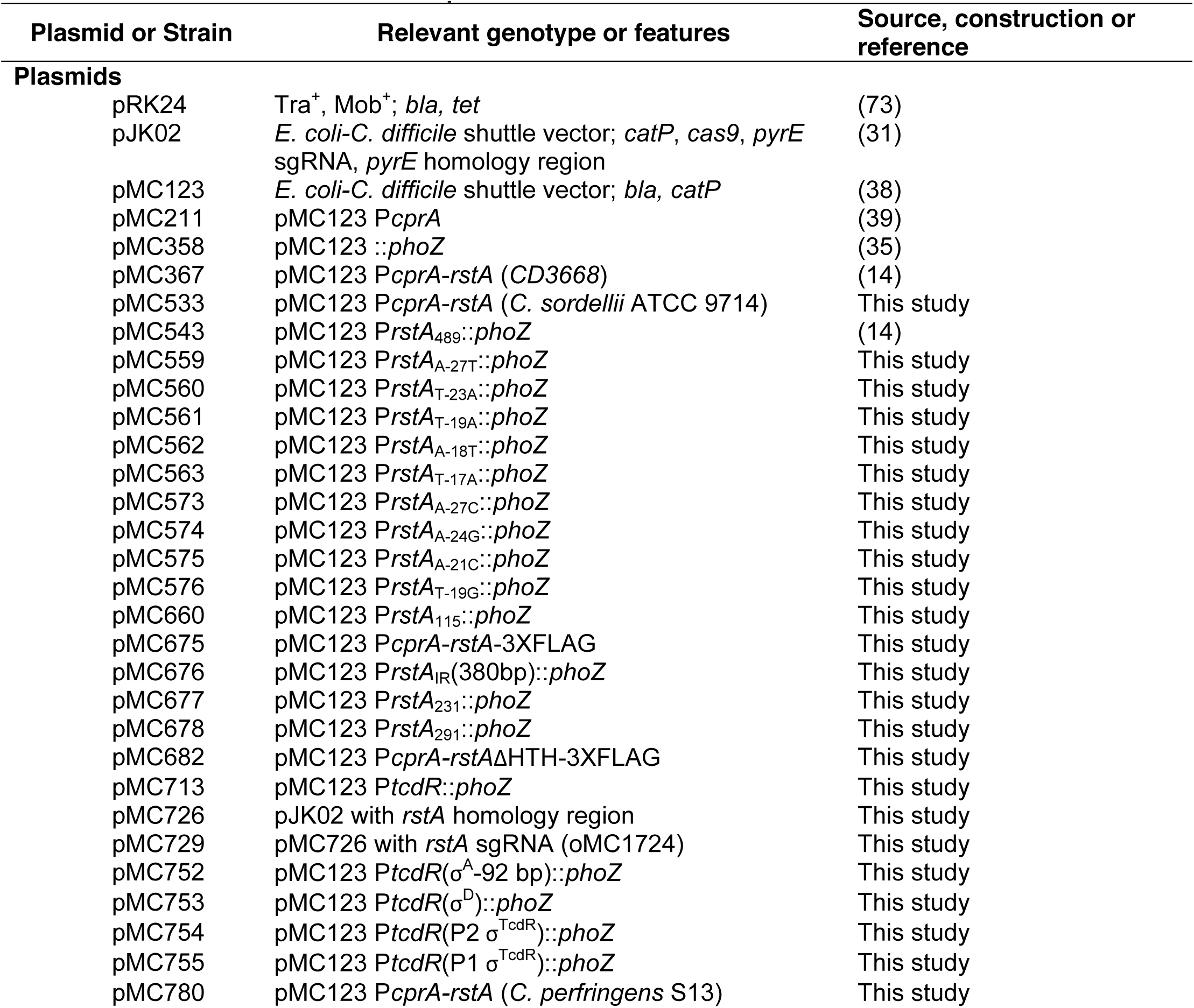
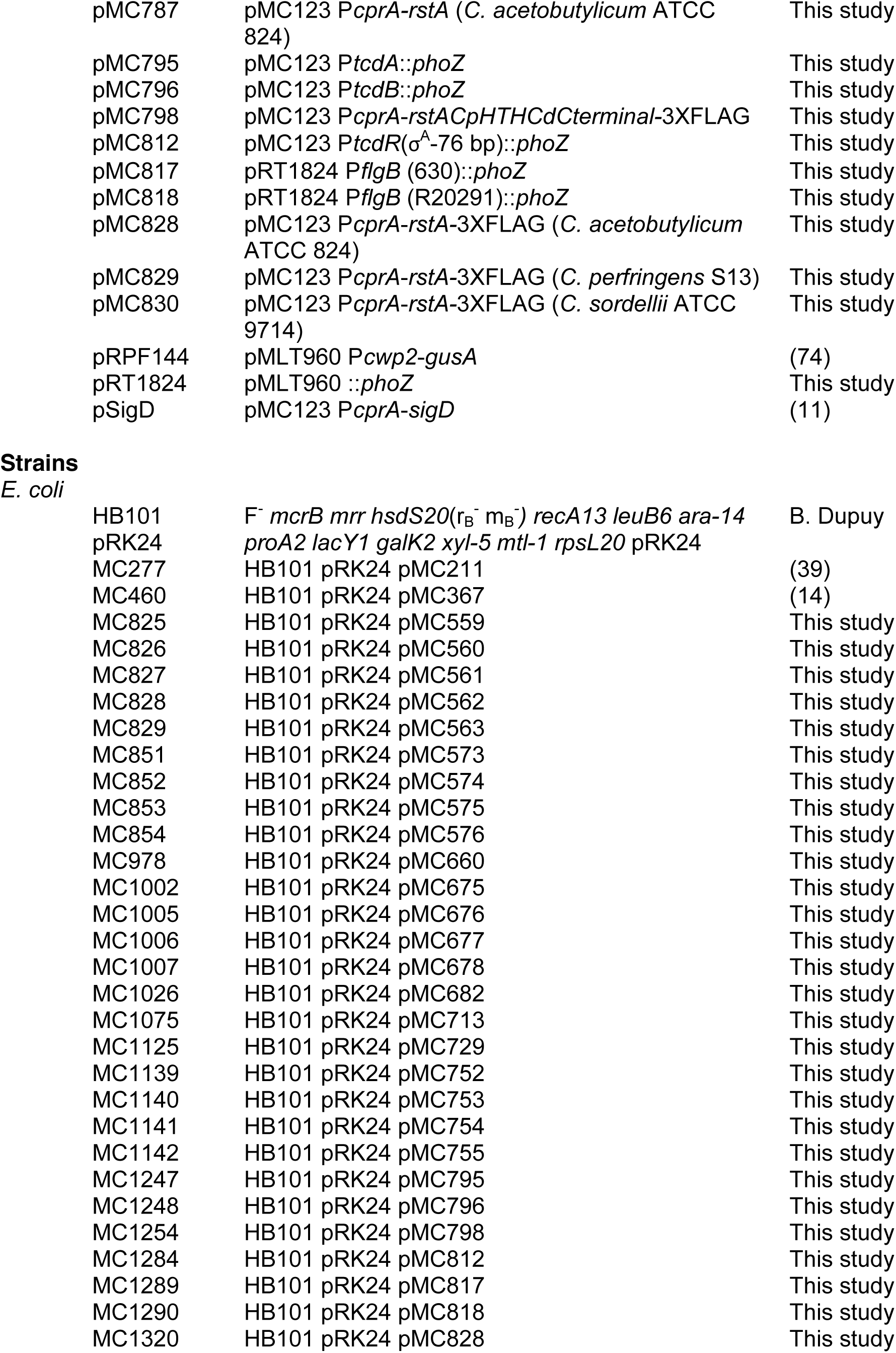
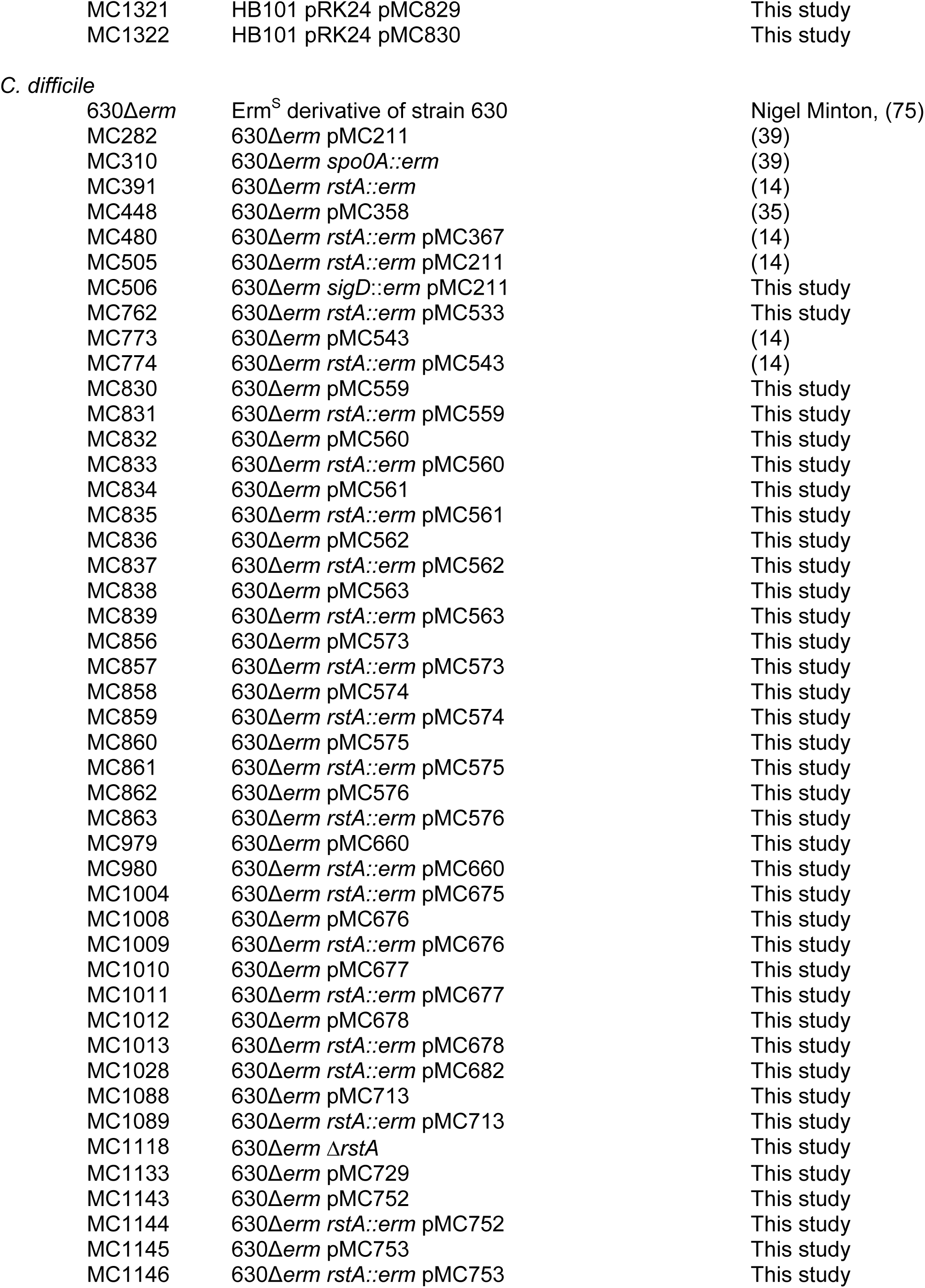
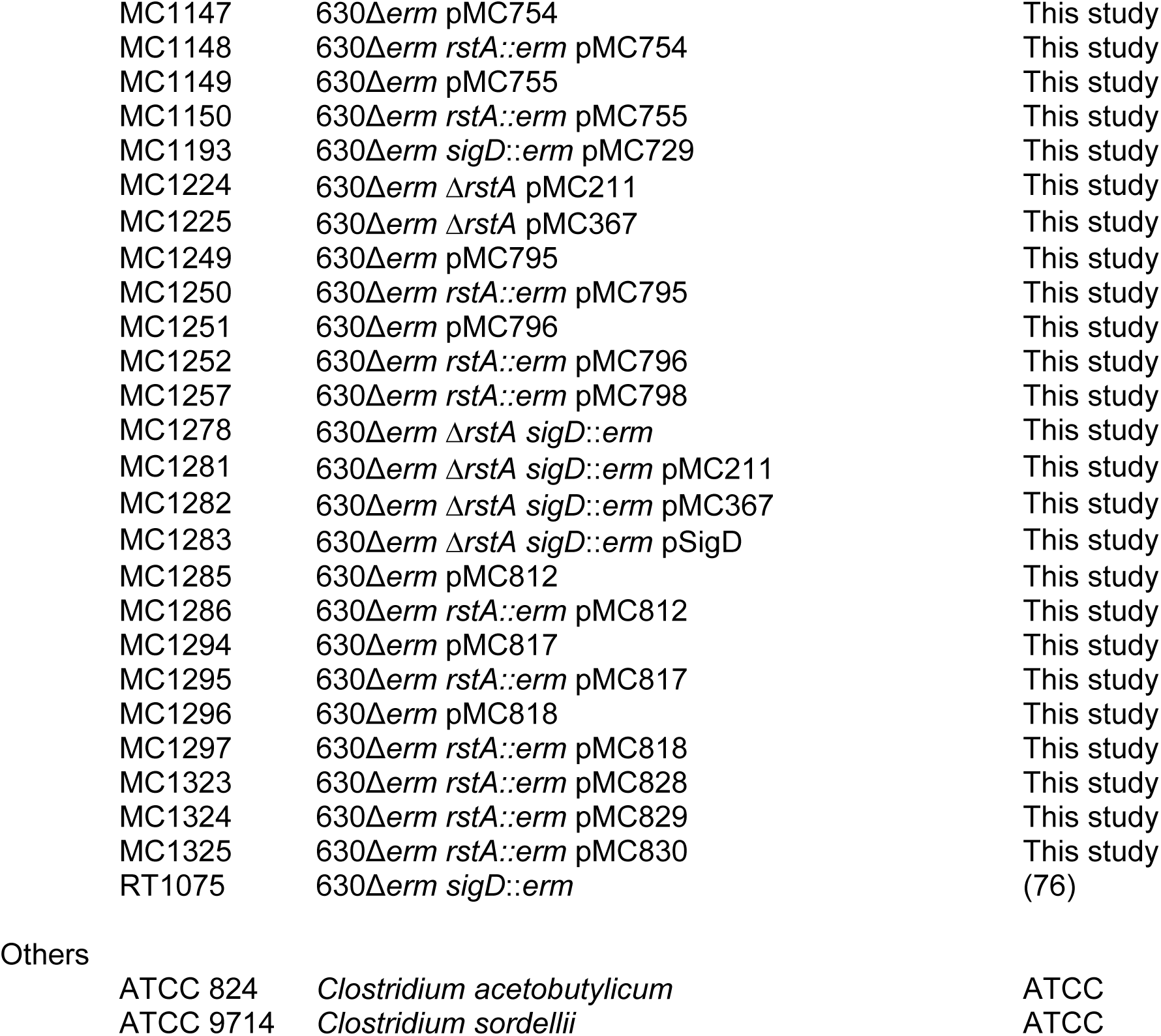
Bacterial Strains and plasmids

**Table S2.**
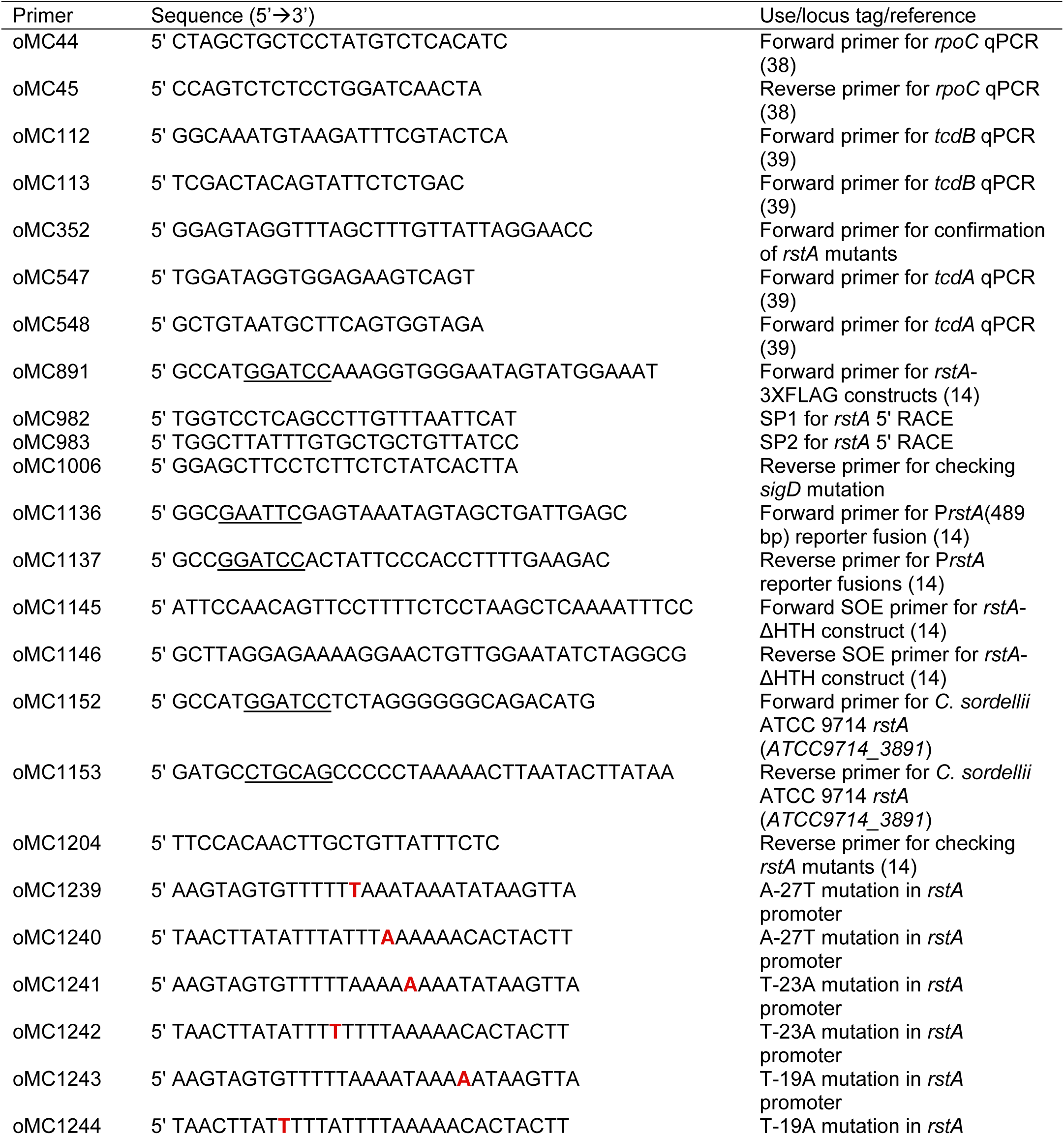
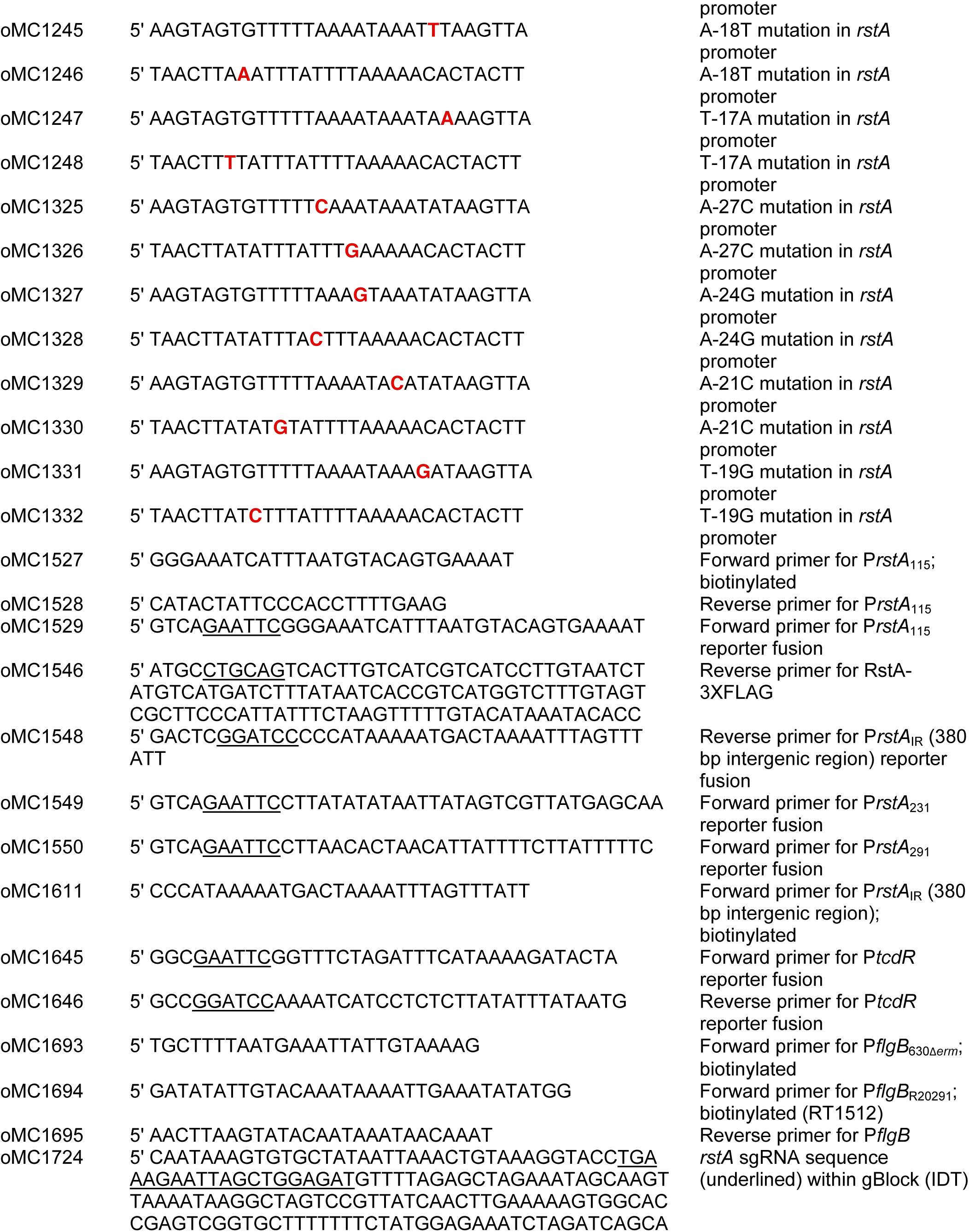
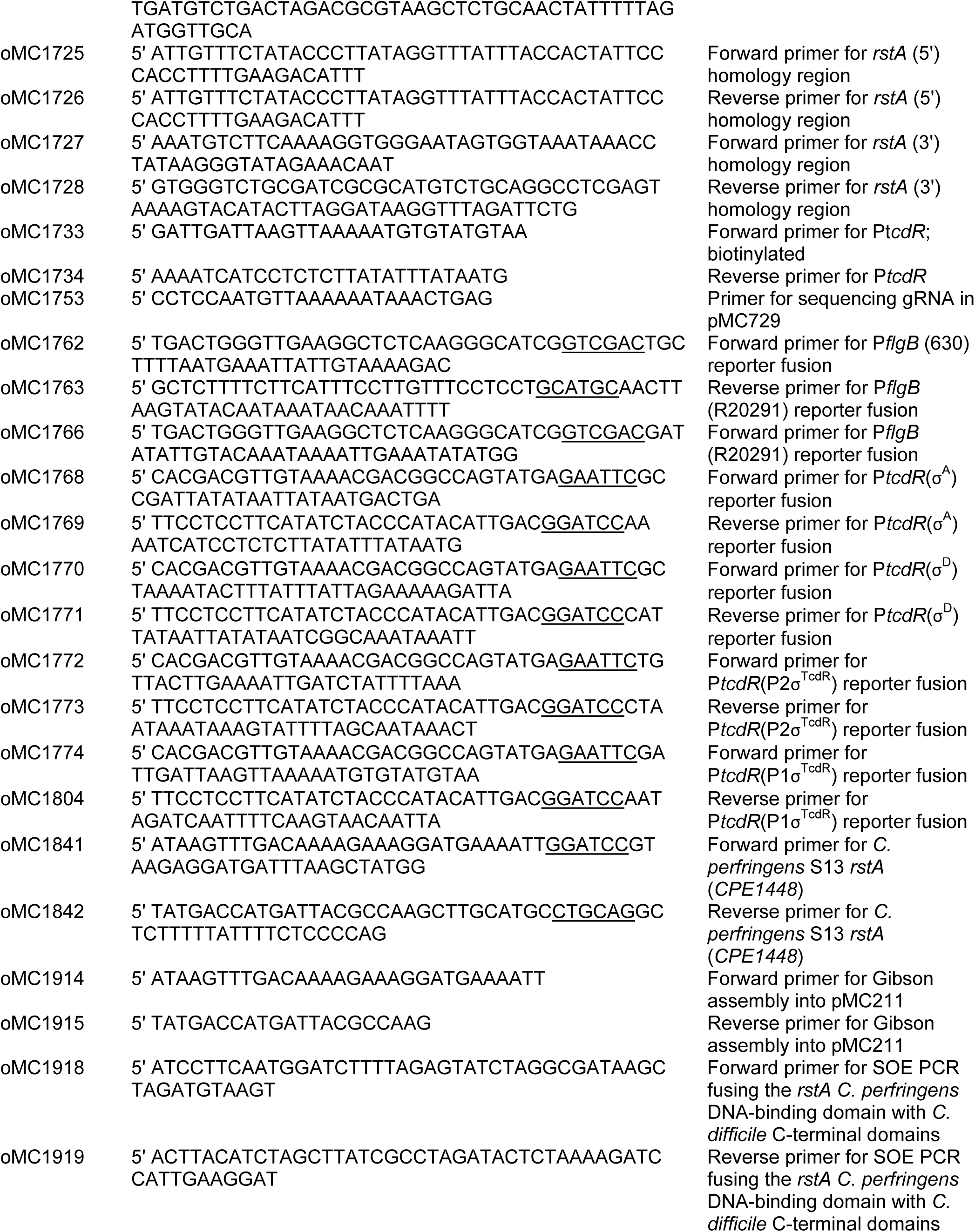
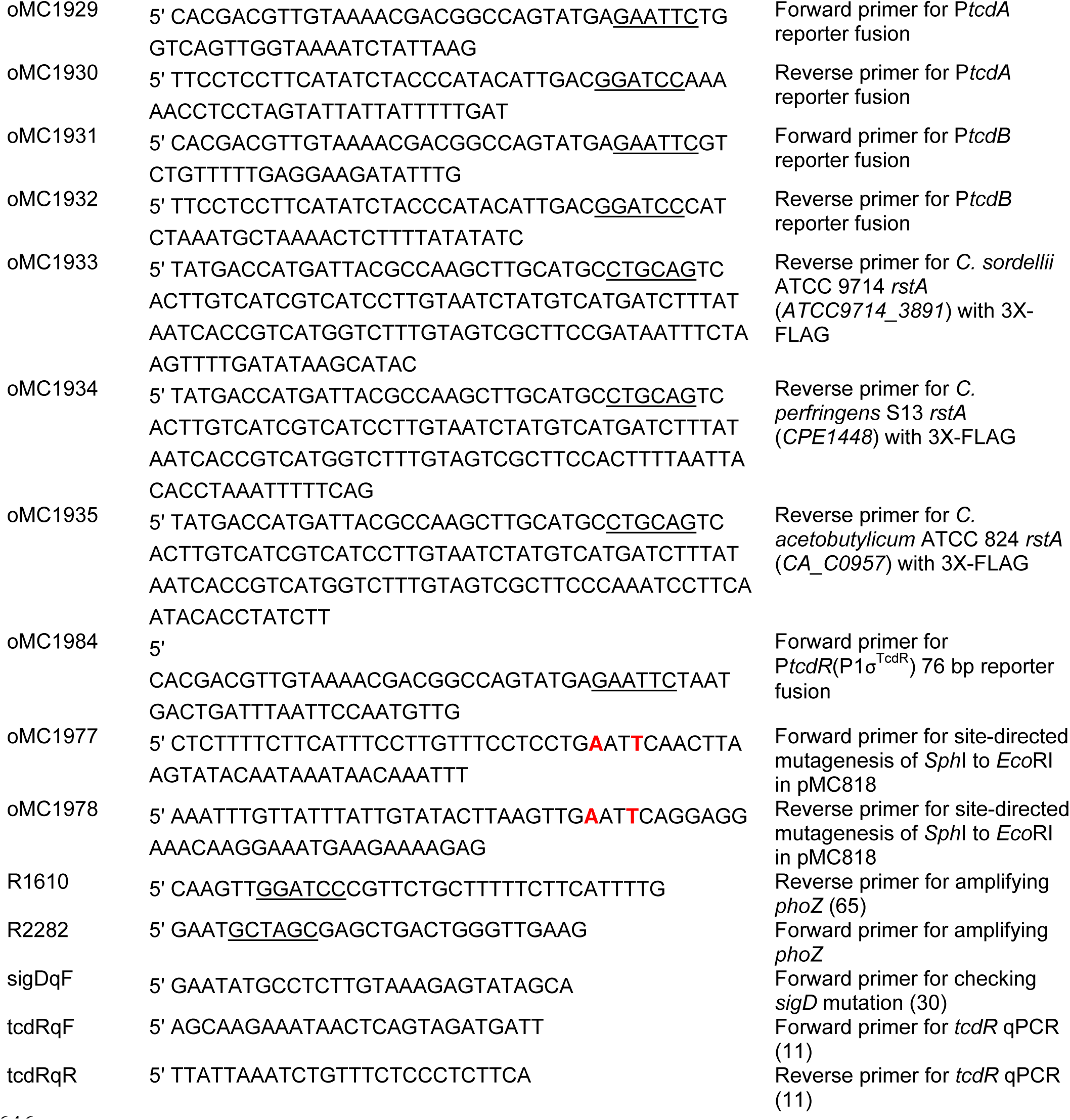
Oligonucleotides. Underlined nucleotides denote the restriction sites used for vector construction. Bolded nucleotides indicate the base mutated within the inverted repeat overlapping the *rstA* promoter.

